# DISTRIBUTION OF LIVING (ROSE BENGAL STAINED) BENTHIC FORAMINIFERA IN COPANO, ARANSAS, AND REDFISH BAYS, CENTRAL TEXAS COAST

**DOI:** 10.1101/2024.12.06.627234

**Authors:** Anna Barrera, Christopher M. Lowery

## Abstract

A combination of human development and climate change have the potential to impact ecosystems in the ecologically, economically, and culturally important bays of the Texas Coastal Bend in the northwestern Gulf of Mexico. To determine the extent to which potential impacts are becoming actual impacts, population studies are necessary. Here, we report populations for living benthic foraminifera from three bays collected in May 2023 and 2024. We find the bays largely dominated by *Ammonia*, a shift from the last complete census carried out by Frances Parker and colleagues in 1951 during a multiyear drought. We see no correlation between predominance facies and salinity within our samples, although we only record a salinity range of 25-30 PSU, a small fraction of the possible range of ∼10-40 PSU in these bays. We do observe a relationship between sediment grain size and predominance facies, with *Ammonia* more common in finer grained environments.

## INTRODUCTION

Estuarine ecosystems in the Gulf of Mexico are being impacted by a suite of changes related to human activity and climate change, including sea level rise, warming temperatures, changing salinity, and degraded water quality. Anthropogenic developments in and around the Texas Coastal Bend add external stress through urban and agricultural run-off, dredging, brine release, and temperature rise. This area has always been prone to large swings in salinity due to changing hydroclimate conditions (Evans et al., 2012), with major implications for salinity-sensitive estuarine species (Parker, 1955; Buzas-Stephens, 2011). Increasing hydrologic extremes are an expected part of climate change (Senevirante et al., 2021) and will in turn make the swings in salinity more severe. Engineering projects are also altering the salinity of these bays. The deepening of Aransas Pass to allow the passage of very large liquefied natural gas carriers to the terminals in the Port of Corpus Christi has increased the volume of water that passes through the inlet with each tide, with impacts on estuarine species (Brown, et al., 2004; Valseth, 2021). Additionally, to meet increasing needs for freshwater, stakeholders in Corpus Christi (including the city, the port, and private industry) are planning to build up to five desalination plants, with some plans calling for the release of brine directly back into bay waters (City of Corpus Christi, 2022). The contract for the first of these plants, in the Corpus Christi Inner Harbor, was awarded to a contractor in October 2024 (City of Corpus Christi, 2024).

Although the modelled salinity increase in the bays from the brine is less than the annual salinity range, this effluent is typically warmer than surrounding water and contains a greater concentration of toxic chemicals than typical seawater (Soliman, et al., 2021).

Tracking ecosystem changes caused by these anthropogenic stressors is urgent, because such data are necessary to plan interventions and policy changes to preserve these important ecosystems. Our knowledge of the effect of climate change and other human activities on coastal ecosystems often only extends as far back as the date of the first observational study at a particular place, limiting our understanding of the extent and duration of change, and of the natural variability of these ecosystems. Benthic foraminifera are sensitive to environmental parameters like salinity, pollution, dissolved oxygen, and nutrient flux, and are commonly used in environmental monitoring studies (e.g., Frontalini and Coccioni, 2011; Martins et al., 2016, 2019; Soukhrie et al., 2017, Sousa et al., 2020). They offer an advantage in paleoecological studies over other larger organisms (e.g., oysters, fish, etc.) because they are ubiquitous in marine environments and can be found in high abundance in small volumes of sediment, making it easy to reconstruct a statistically significant species-level population.

Benthic foraminifera have been previously studied in Aransas Bay and Copano Bay, in what is today the Mission Aransas National Estuarine Research Reserve. These bays have fewer direct human impacts than adjacent Corpus Christi Bay (especially in terms of dredging), making them ideal for understanding the benthic foraminiferal response to environmental change. The total core top assemblage of benthic foraminifera was reported for Aransas and Copano bays by Parker et al. (1953) from samples collected in 1951. These data include tests which were dead at the time of collection and thus represent a multiyear average population. Living (Rose Bengal stained) foraminifera were reported for Aransas Bay by Phleger (1956) and Copano Bay by Buzas-Stephens et al. (2018) from samples collected in 1954 and 2006, respectively. Phleger’s sample volumes were small, however, and nearly half of his Aransas Bay samples contained fewer than 100 specimens.

Here, we report living (Rose Bengal stained) benthic foraminifera collected in May 2023 and 2024 in Redfish, Aransas, and Copano Bays (Figure 1), all within the Mission-Aransas National Estuarine Research Reserve. As far as we are aware, our paper is the first quantitative report on living benthic foraminifera in Redfish Bay, although Ladd et al. (1957) reports qualitative abundances of benthic foraminifera from Redfish Bay based on observations in 1940. These populations thus mark a modern baseline against which to measure future change and an opportunity to determine how the community has changed in the past decades. Poag (2015) used the work of Parker et al. (1953) to define his benthic foraminiferal predominance facies in these bays; in 1951, most of Aransas Bay was characterized by a roughly even split of *Ammonia* and *Elphidium* species and was thus mapped as *Ammonia/Elphidium* predominance facies. In this work we set out to test whether that predominance facies measured in the 1950s still typifies Aransas Bay. Interestingly, we found that, in May 2023 and 2024, Aransas Bay assemblages were nearly entirely *Ammonia*.

**Figure 1.**
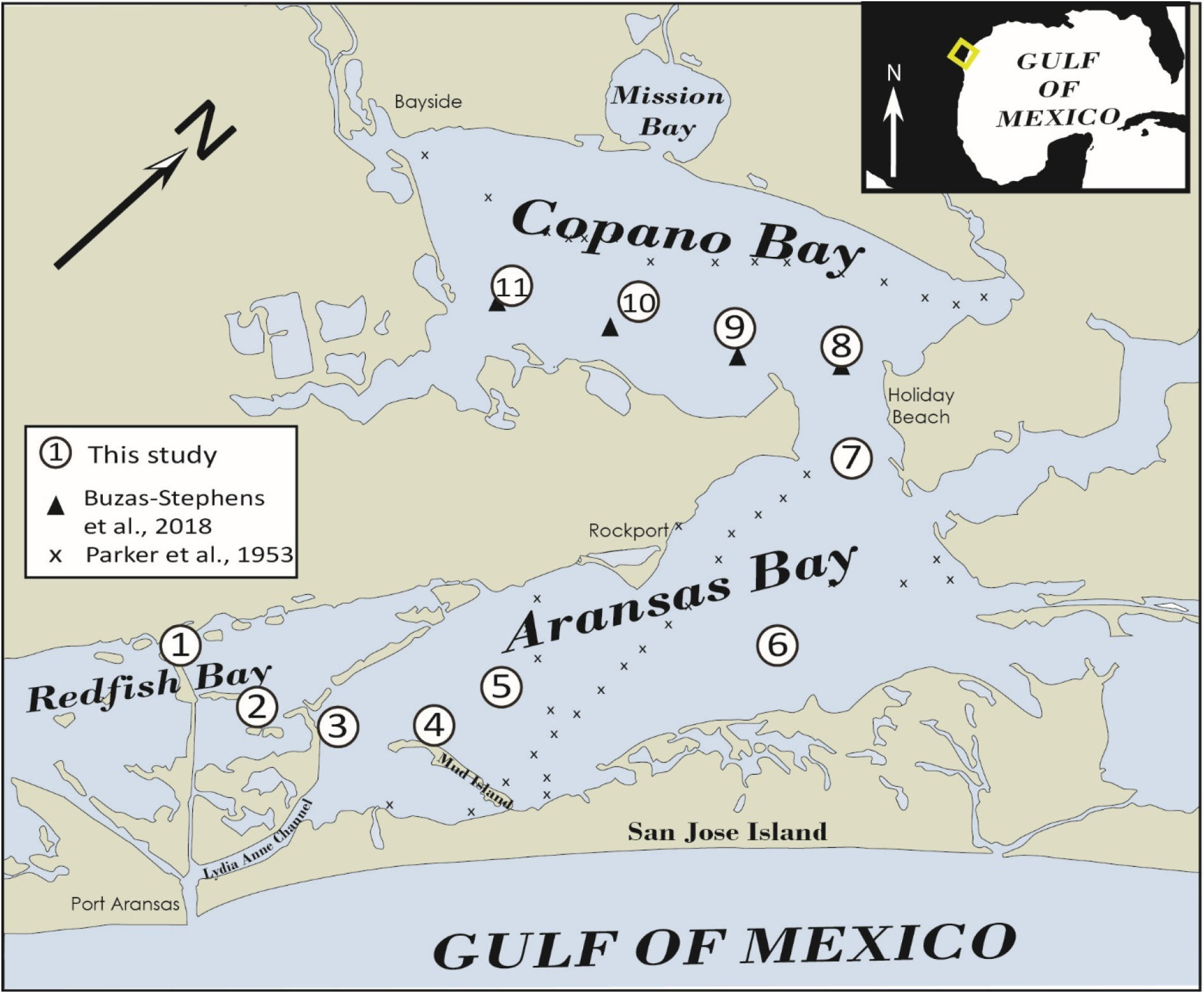
Location map showing the three bays studied and sample locations. Map traced from the “Oceans” basemap on the NOAA arcgis map viewer (NOAA 2024).

## METHODS

### Sample Collection

Samples analyzed here were collected during two consecutive years of the University of Texas at Austin’s Marine Geology and Geophysics Field Course, an upper level course taught each May, with the field component based in Port Aransas, TX. Samples were collected using a Ponar grab sampler deployed from the R/V *Curt Johnson*. A total of seventeen samples were collected from the eleven stations, including nine samples May 10-13, 2023, and eight samples May 15-18, 2024 (Table 1). Some stations did not have a successful sample retrieval in both years. In Copano Bay this was the result of oysters in the sampler (we attempted to solve this problem by offsetting a few dozen meters, but merely found more oysters); in Aransas Bay the cause of the sampler repeatedly coming up empty was less obvious but was likely due to some sort of firm substrate. At each station, surface salinity was measured with an optical refractometer and recorded. The optical refractometer displays salinity in units of per mille (‰), although here we report it in the equivalent international standard Practical Salinity Unit (PSU). Location was recorded using a Garmin inReach handheld GPS, which has a horizontal resolution of 5-10 m under normal conditions.

**Table 1.**
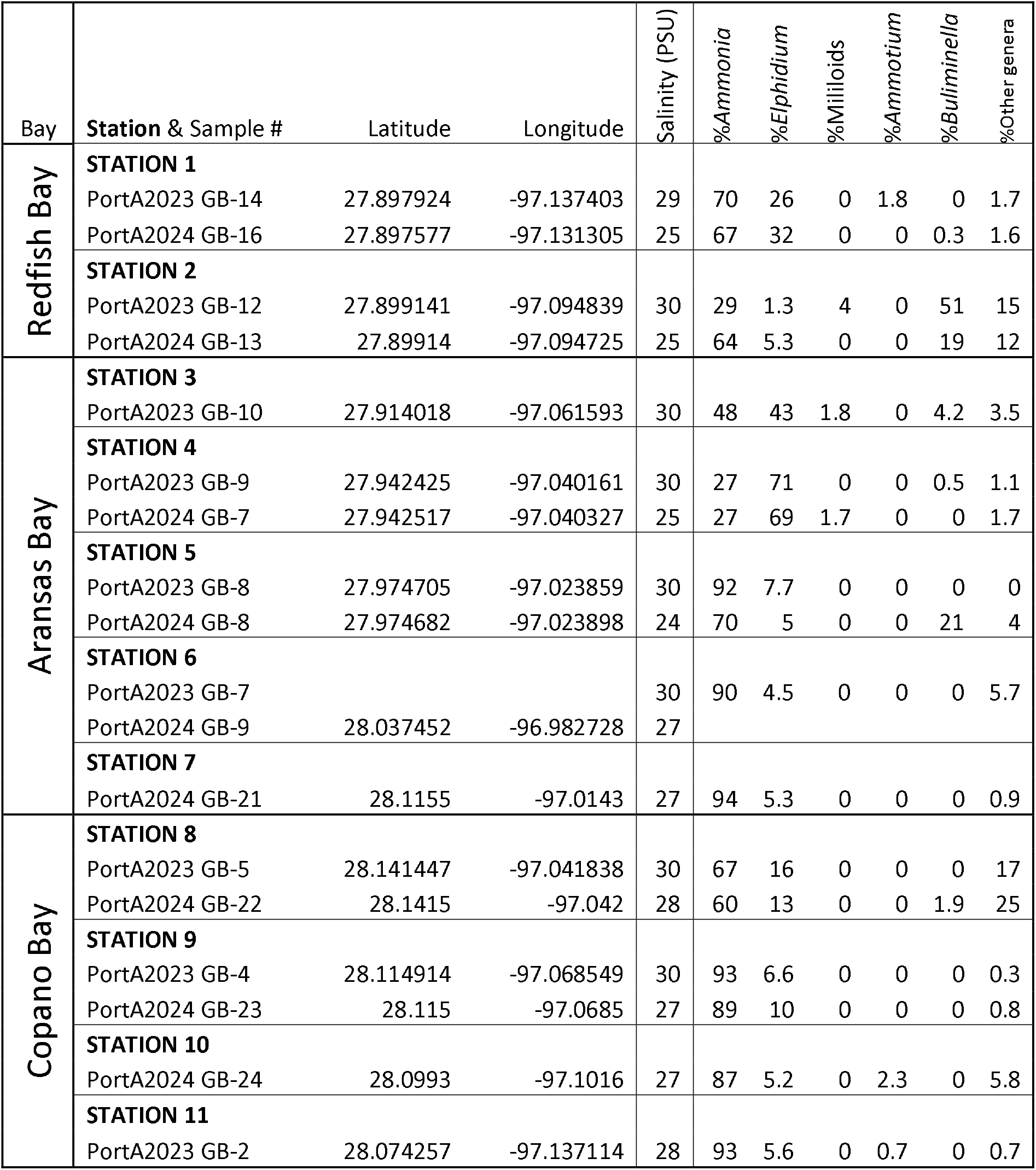
Sample names, sample locations, measured salinity, and percentage abundance of common living (stained) foraminifera found in each. Individual samples are labelled by campaign, year, sampler type (GB = grab sampler), individual sample number. Station numbers were assigned post-hoc based on the samples selected for this study. See supplemental data for a more detailed population table.

### Sample Processing and Analysis

Immediately after retrieval, the upper centimeter of sediment was scraped off the top of the grab sample, placed in a Nalgene bottle, and covered with a solution of 2% Rose Bengal, 4% formalin (a fixant), and sodium hexametaphosphate (a buffer). This mixture of sample and solution was given a good swirl to break up the mud, then placed in a cooler for at least 24 hours. The sample then was washed over a 63 micron sieve and dried in an oven at 75°C. Back in the lab, the sample was split in a microsplitter to obtain a manageable amount, and stained foraminifera were identified and picked under a Zeiss Stemi binocular microscope. We aimed for at least 300 stained specimens per sample (e.g., Patterson and Fishbein, 1989) but many samples did not yield this many; all samples except for one contained more than 100. “Stained” specimens were defined as specimens in which at least one chamber was fully stained pink; specimens with surface staining, speckled staining inside one or more chambers, or which were lightly tinted pink were not included in the counts. Since we were picking samples dry, it was necessary to wet opaque tests (miliolids and agglutinated taxa) with the brush to determine staining.

Specimens were identified using species lists from Parker et al., 1953 updated to modern species concepts based on the work of Poag (2015). Many estuarine taxa, *Ammonia* in particular, have a rich history of species concepts that have been synonymized, split, relegated to subspecies, and then resurrected by different workers (Haynes, 1992, lamented “nomenclatural chaos” when writing about *Ammonia*). Acknowledging this, we follow the lead of Poag (2015), whose book on Gulf Coast foraminiferal assemblages names the common species of this genus *Ammonia parkinsoniana*, and recognizes two ecophenotypes as subspecies, a large, thick-walled “forma *typica*” and a small, thin walled “forma *tepida*.” In our data, we do not differentiate between these ecophenotypes and simply report *A. parkinsoniana*, the only species of the genus *Ammonia* which we identified in our samples. Because we are primarily concerned with testing the predominance facies of Poag (2015), which are focused on generic abundances, we focused on counts at the genus level and report them as such in Table 1. To account for sampling error, confidence intervals for each genus were calculated using the method of Patterson and Fishbein (1989). Predominance Facies were defined by identifying the dominant genus in each sample. Because we did not sample a consistent volume of sediment, we do calculate total foraminifera.

Grain size analysis was conducted the same day as sampling by the field course students using the pipette-sieve method, where the gravel (>2000 microns) and sand (2000-63 microns) size fractions were measured directly over sieves, and the silt (63-4 microns) and clay (<4 microns) size fractions were calculated from aliquots taken from the <63 micron fraction after settling in a 1000 ml graduated cylinder for 57 seconds (silt+clay fraction) and 2 hours and 3 minutes (clay fraction).

## RESULTS

### Salinity

Average salinity across all stations was about 4 PSU higher in May 2024 than May 2023. In May 2023, salinity at our stations averaged 25.0 PSU in Redfish Bay, 25.3 PSU in Aransas Bay, and 27.2 PSU in Copano Bay. In May 2024, salinity at our stations averaged 29.6 PSU in Redfish Bay, 30.0 PSU in Aransas Bay, and 29.3 PSU in Copano Bay. The Aransas River is the main source of freshwater into these bays, along with the smaller Mission River, which covers an adjacent catchment. For the month prior to sample collection in 2023, discharge of the Aransas River at the USGS stream gauge near Skidmore, TX averaged 0.12 cubic meters per second. In 2024, it averaged 0.08 cubic meters per second during the same period.

### Grain Size

Grain size varied throughout the study area, with samples from Redfish Bay dominated by sand and samples from Aransas and Copano Bays dominated by mud (defined here as silt + clay). The gravel size fraction, comprised of whole and broken bivalve shells, was variable from station to station. Grain size measurements for each station are included in the supplemental material. Grain size varied sightly from year to year at every station, which is likely reflective of the heterogeneity of the seafloor in bay environments even over short distances. Redfish Bay samples averaged 71% sand and 20% mud over both sampling campaigns. Aransas Bay samples averaged 41% sand and 54% mud, although there was more sand in samples in the southwestern part of the bay, near Mud Island (a poorly named sandy spit on the back of San Jose Island) and near the mouth of Lydia Anne Channel. Samples in Copano Bay averaged 46% sand and 54% mud. There, the sandiest samples were from Station 8, just inside the entrance to Copano Bay and perhaps reflective of the channeling of tidal energy at the mouth of the bay.

### Foraminifera

The nine samples collected in 2023 contain a range of 26 to 305 individuals per sample and 14 genera total. The eight samples collected in 2024 contain a range of 103 to 310 individuals per sample and 13 genera were identified. Simplified count data are shown in Table 1; full counts including rare taxa are included as a supplemental table. Overall, all three bays were dominated by *Ammonia*, with *Elphidium* becoming more common in the southwestern part of Aransas Bay, near Lydia Anne Channel (which leads directly to the Aransas Pass inlet), and a single sample in a seagrass bed in Redfish Bay dominated by *Buliminella gracilis*.

In Redfish Bay in 2023, the predominant genus was *Ammonia*, which made up 70.2% of the assemblage at Station 1 and 29.3% at Station 2. (Table 2). The second most common genus was *Buliminella*, which dominated Station 2 with 50.7%. This station was in a seagrass bed near Hog Island, and the grab sampler came up with numerous blades of grass in its jaws. *Elphidium* made up 26.3% of Station 1 and just 1.3% at Station 2. Samples from 2024 in Redfish Bay were similar overall, although Station 2 had far less *Buliminella* (18.8%) and did not contain any seagrass (this more likely reflects the spotty nature of seagrass cover than any major shift in the ecosystem). Instead, *Ammonia* made up 64.3% of Station 2, similar to the 66.5% of Station 1.

In central and northern Aransas Bay, *Ammonia* was the dominant genus in both 2023 and 2024, comprising up to 93.8% of the assemblage (Figure 2). Most of the rest of these samples are rounded out with single digit percent abundances of *Elphidium*, with a few *Haynesina* present in low abundance and a significant assemblage of *Buliminella* (21%) at Station 5 in 2024 (this taxon was completely absent there in 2023). However, *Elphidium* becomes increasingly abundant in the southwestern part of Aransas Bay, dominating Station 4 near Mud Island with ∼70% both years and comprising about equal parts, along with *Ammonia* (42.8% and 47.7%, respectively), of Station 3 north of the tidally dominated Lydia Anne Channel, which connects directly to the Aransas Pass inlet. We map the area around these two stations as mixed *Ammonia/Elphidium* facies on Figure 2, but it is likely this area is undersampled and may exhibit more variability than we can see from the data we present here. It is interesting to note that at the time of sampling these two stations had nearly identical salinities to the other stations in Aransas Bay (and Copano Bay, for that matter).

**Figure 2.**
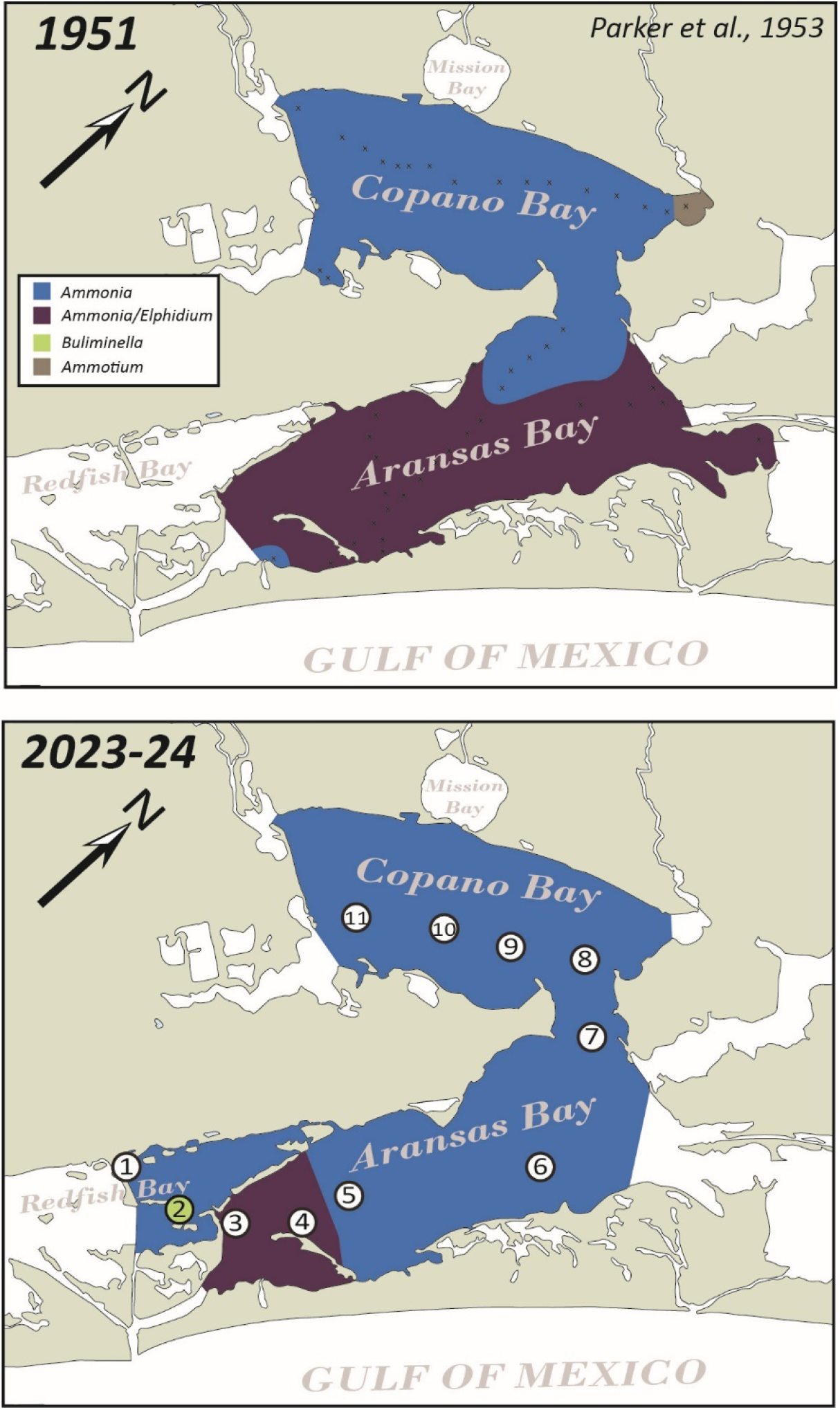
Predominance facies maps showing the dominant genus or genera across the study area in 1951 (Parker et al., 1953) and 2023-2024 (this study). Predominance facies did not change from 2023 to 2024, with the exception of station 2 (taken in a seagrass bed), which was dominated by *Buliminella gracilis* in 2023 (when the sample contained numerous blades of seagrass) and *Ammonia parkinsoniana* in 2024 (when the sample contained no seagrass).

Copano Bay was dominated by *Ammonia* both years (Figure 2), although the degree of its dominance varied from east to west. At the eastern most Copano Bay Station, Station 8, *Ammonia* comprised 67.3% of the population in 2023 and 60.2% in 2024. Other common taxa included *Elphidium* (15.7% and 12.6%, respectively), followed by *Bolivina* (10.6% and 10.7%) and various agglutinated taxa. Further west, in Stations 9-11, *Ammonia* was present at similar levels to Aransas Bay, representing 86.7 to 93.1% of the assemblage.

## DISCUSSION

### Difference in Assemblages Since the 1950s

The most striking feature of the benthic foraminiferal population in 2023 and 2024 is how different it is from the last census of benthic foraminifera in these bays in 1951 (Parker et al., 1953) (Figure 2). In August 1951 Frances Parker, Fred Phleger, and Jean Pierson took 37 samples in Aransas Bay and 16 samples in Copano Bay using a small coring tube and examined the upper 1 cm of sediment for total (living + dead) foraminifera. In both Aransas and Copano Bays they found assemblages with a much higher proportion of *Elphidium* than was observed in 2023 and 2024. In Copano Bay, although *Ammonia* still dominated the assemblage, *Elphidium* averaged 19% of the population (compared to 9% in 2023-24) (Figure 2). In Aransas Bay, *Ammonia* and *Elphidium* were roughly equal, representing 45% and 38% of the mean assemblage, respectively. Today, *Elphidium* is common in the far southwestern part of Aransas Bay, but is only present in single digit percent abundances through most of the bay (Figure 2).

In 2006, Pamela Buzas-Stephens and colleagues sampled four stations in Copano Bay, and found living assemblages that are distinctly similar to ours, with three of the four stations similarly dominated by *Ammonia* (average 78% of the population) and the fourth nearly evenly split between *Ammonia* and the agglutinated genus *Ammotium* (Buzas-Stephens et al., 2018), which is a common component of populations proximal to fresh water inputs (Poag, 2015).

Why is there such a large difference between populations in 1951 and 2006/2023-24? The most likely cause is salinity. Salinity varies from ∼10 PSU to ∼40 PSU on multiyear scales in Aransas and Copano Bays (Evans et al., 2012). Sampling in 1951 occurred in the middle of an extended drought that lasted from 1948 to 1953 (Parker, 1955). Salinity at the time of sampling was very high in the bays, ranging up to 42 PSU (compared to 36 PSU in the open Gulf of Mexico at the time; Parker et al., 1953). This extended drought is associated with major changes in the macroinvertebrate community of the Coastal Bend estuaries and lagoons as open ocean taxa, better adapted to higher salinities, displaced estuarine species which were not (Parker et al., 1955). Echinoderms and corals invaded Aransas Bay as far north as Rockport, and Gulf Crabs became more common in the bays (Parker, 1955). The shrimp catch dropped from 1949 to 1951 (Hildebrand and Gunter, 1953) and production of the common oyster (*Crassotrea virginica*) was reduced (Parker, 1955). Parker (1955) reports that oysters from 1948-1950 were “thin, flabby, and tasteless” and by 1952 oysters were dying in large numbers in Copano Bay. Oyster reefs became dominated by the more salinity-tolerant small oyster *Ostrea equistris* (Parker, 1955).

Against this backdrop of major changes in ecologically and economically important macroinvertebrate species, Parker et al. (1953) observed a high occurrence of *Elphidium*, which is commonly found in the open gulf and, in other Texas bays, dominates the most distal bay waters, near the inlet (Poag, 2015 and references therein). *Ammonia*, meanwhile, is also commonly found in the open Gulf and Texas bays, but within bays it occupies a more proximal position relative to the bayhead delta compared to *Elphidium* (Poag, 2015), perhaps suggesting a tolerance for lower salinity waters. It is thus reasonable to assume that a long period of drought and increased salinity caused an increase in the number of *Elphidium* in Aransas and Copano Bays in the 1950s, whereas more normal precipitation and lower salinities in 2006 and 2023-24 resulted in a higher dominance of the more brackish-tolerant *Ammonia*.

The data of Parker et al. (1953) are the total core-top assemblages, which includes both living and dead foraminifera, and so comprise a longer-term average than a dataset composed of only foraminifera which were living at the time of sampling. It is therefore possible that the impact of the drought on the benthic foraminiferal community in Aransas Bay was even more severe than is suggested by these populations. On the other hand, it is also possible that the normal long term average population in Aransas Bay tends to have more *Elphidium* in it, and our living foraminiferal data are shorter snapshots of time intervals more dominated by *Ammonia*. A longer time series of living foraminifera from the modern bay will be necessary to test this.

Fred Phelger collected living (Rose Bengal stained) foraminifera in Aransas Bay in June 1954, immediately following the drought (Phleger, 1956). Salinities in Aransas Bay ranged from 31-36 PSU, roughly intermediate between those of the drought ∼40 PSU and those recorded during our sampling in 2024 (∼30 PSU). Of the 10 samples he collected, five contained fewer than 100 specimens. The other five contained assemblages dominated by *Ammonia* but with a larger proportion of *Elphidium* (up to 30%) than we see in our samples today (Phleger, 1956). The sample with the highest proportion of *Elphidium* was directly adjacent to Mud Island, near our Station 4 (which also contained the maximum *Elphidium* we observed). Overall, these samples represent an intermediate stage between the high-salinity drought conditions sampled by Parker et al. (1953) and the moderate salinity conditions we observed in 2023-24.

### Controls on Modern Species Distribution

While a major sustained shift in salinity is an intuitive explanation for the differences from the 1950s to 2000s, the direct ecosystem impact of fluctuating salinity conditions below the level of a severe multi-year drought is not clear. In our dataset we see an increased percentage of *Elphidium* in the southwestern part of Aransas Bay, near the connection to Lydia Anne Channel, Aransas Pass, and the open Gulf of Mexico, suggesting a more marine affinity for this genus compared with *Ammonia*, as we might expect if its distribution is driven by salinity (as is suggested by the predominance facies maps across the region by Poag (2015). However, when we compare the percentage of *Elphidium* or *Ammonia* in our samples with the seawater salinity measured at the time of collection, we see absolutely no correlation (Figure 3; the R^2^ value for those lines is 0.0002). Salinity in these bays can fluctuate on a short term basis, and it is possible that such fluctuations obscure longer salinity trends related to distance from the inlet or from freshwater input that control foraminiferal distributions. Or, there could be some other physiographic parameter or set of parameters (nutrient supply, seafloor oxygen content, turbidity, temperature, pH, etc.) that we did not measure that may play a role.

**Figure 3.**
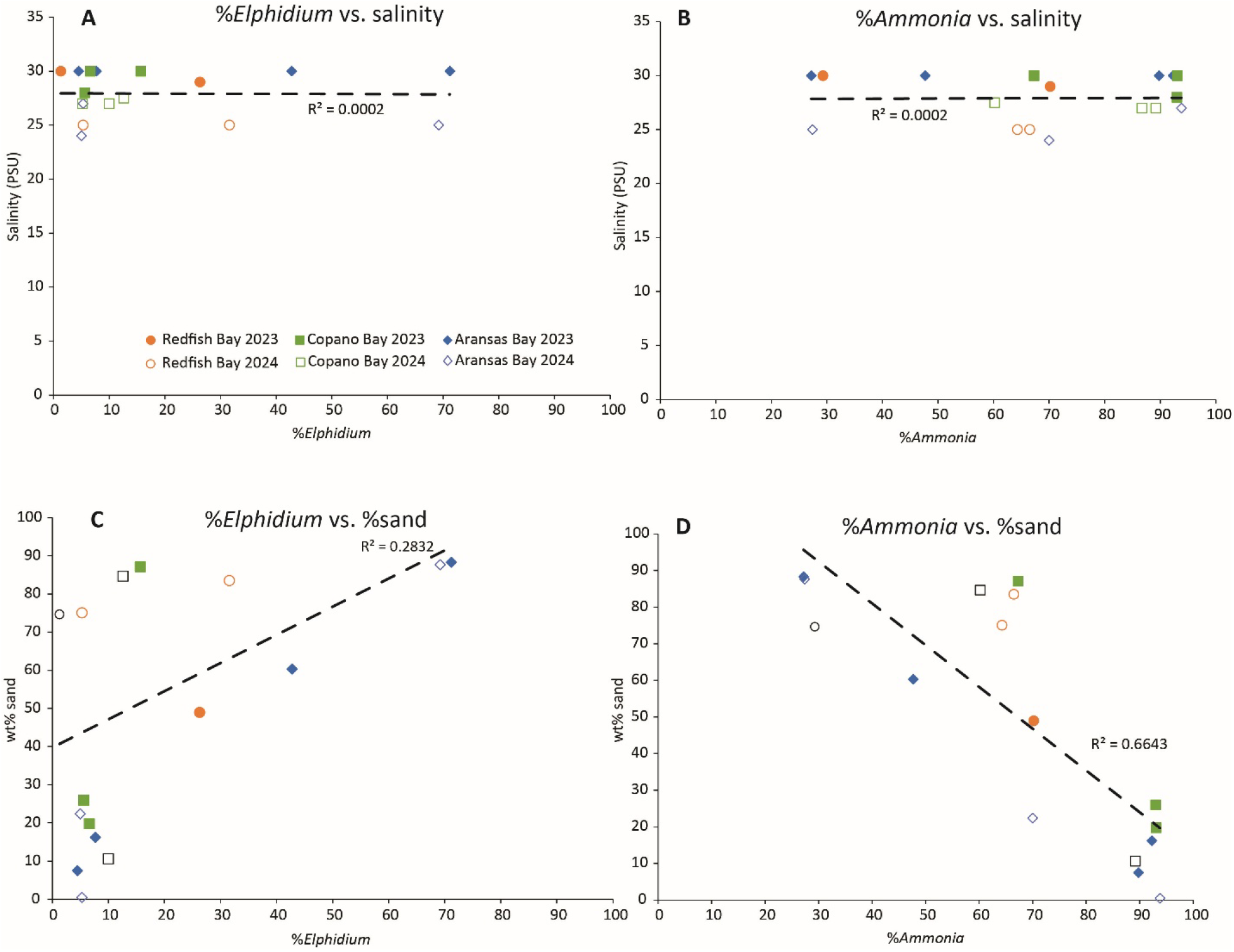
Crossplots of the two most common genera against salinity (A-B) and %sand (C-D) in all three bays for both years. Circles represent samples from Redfish Bay, squares Copano Bay, and diamonds Aransas Bay, with filled symbols for 2023 and open symbols for 2024. Within our dataset, salinity shows no correlation whatsoever with A) %*Elphidium* or B) %*Ammonia*. C) %*Elphidium* shows a weak positive correlation with %sand, while D) %*Ammonia* shows a stronger negative correlation with %sand.

There is one other parameter important that we did measure during the field course, and that is grain size. In non-polluted coastal environments, sediment type is an important factor in the distribution of foraminiferal biofacies (e.g., Armynot du Châtelet et al., 2009; Celia Magno et al., 2012). In stark contrast to salinity, the percentage of sand in each sample negatively correlates with the relative abundance of *Ammonia* (Figure 3D). *Ammonia parkinsoniana* is commonly associated with soft, muddy substrates (Armynot du Châtelet, 2009; Suokhrie et al., 2017). Because of this, it is also commonly found at polluted sites (as pollutants tend to settle on the bottom in lower energy, muddier areas) and sites with high sedimentary organic carbon content (TOC) (e.g., Armynot du Châtelet et al., 2009; Suokhrie et al., 2017). We do not have trace metal or TOC data for these sites, and unfortunately the NERR water quality monitoring stations in Copano Bay and Aransas Bay stopped recording nutrient and chlorophyll data at the end of 2022. But we expect that the muddier sediments (which often had the distinct smell of decaying organic matter) are elevated in TOC.

Our work here demonstrates that the benthic foraminiferal population of these bays is not stable and can change significantly over time periods of at least several decades, if not faster. Predominance facies defined in the literature for particular bodies of water, such as those presented by Poag (2015), are likely more ephemeral snapshots of conditions at the time of collection than stable environments which can be used to guide paleoecological interpretation. Continued sampling of these environments is essential to determine their short- and long-term variability (see, for example, the monthly timeseries of Buzas et al., 1977). We have only taken samples across a small portion of the known salinity range in these bays, which ranges from <10 to >40 PSU, and it is possible that a relationship between salinity and dominant taxa may become clear with greater differences in salinity.

## CONCLUSIONS

Living benthic foraminifera collected in 2023 and 2024 from Copano, Aransas, and Redfish Bays reveal a foraminiferal ecosystem dominated by *Ammonia*, with increased *Elphidium* in southwestern Aransas Bay, closest to Lydia Anne Channel and the open Gulf of Mexico, and a dominance of *Buliminella gracilis* in a single sample from a seagrass patch in Redfish Bay. This is different than the population found in these same waters during an extreme drought by Parker et al. (1953), which had a much higher proportion of the more marine-affinity genus *Elphidium*, which those authors attributed to the high salinity conditions in the bays at the time of sampling. It is not entirely clear that salinity is the cause of the change observed since the mid-century, although we suspect it is the most likely. All the published foraminiferal data come from the saltier part of the total possible salinity range of these bays, and more samples must be taken across lower salinities to gain the whole picture. Given ongoing anthropogenic threats to these habitats, a regular census of the benthic foraminiferal population would provide an important way to monitor ecosystem health. Determining the relationship between benthic foraminifera and salinity and other environmental variables in the modern bays will also provide a tool to investigate Holocene sediments and identify high salinity and high nutrient flux intervals in the past. This will allow us to determine the recurrence interval for intense droughts impacting estuarine salinity in the Coastal Bend like that of 1948-1953, and understand the natural variability in these ecosystems, in order to determine if anthropogenic impacts are pushing these bays outside their natural range. Ongoing modern observations at different parts of the salinity cycle are an essential pre-requisite for this. Moreover, these data highlight the need to measure a wide number of environmental variables when collecting living foraminifera.

## Supporting information

supplemental tables of species counts and grain size

## ACKNOWLEDGEMENTS

We are grateful to Captain Frank Ernst of the *R/V Curt Johnson* for cheerfully ferrying us around these bays, and the students of the 2023 and 2024 Marine Geology and Geophysics Field Course for their work collecting these samples. We also thank Carson Miller, Patty Standring, Lorna Kearns, and Adam Woodhouse for their assistance in taking and washing samples, and Marcy Davis, Dan Duncan, Sean Gulick, John Goff, Dallas Sherman, and Jingxuan Wei for logistical help in the field.

## Notes

### Competing Interest Statement

The authors have declared no competing interest.

### Summary of Updates

This manuscript has been revised to reflect peer reviews of the original manuscript in JFR. Main changes include more discussion of the usage of genus-level identification and its limitations, more detailed description of the grain size results (and grain size added to supplemental data), revision to the color scheme in Figure 2, addition of more data to Figure 3, and general revision of the text to improve clarity.

